# Next-generation sequencing analysis with a population-specific reference genome

**DOI:** 10.1101/2024.03.07.584017

**Authors:** Tomohisa Suzuki, Kota Ninomiya, Takamitsu Funayama, Yasunobu Okamura, Shu Tadaka, the Tohoku Medical Megabank Project Study Group, Kengo Kinoshita, Masayuki Yamamoto, Shigeo Kure, Atsuo Kikuchi, Gen Tamiya, Jun Takayama

## Abstract

Next-generation sequencing (NGS) has become widely available and is routinely used in basic research and clinical practice. The reference genome sequence is an essential resource for NGS analysis, and several population-specific reference genomes have recently been constructed to provide a choice to deal with the vast genetic diversity of human samples. However, resources supporting population-specific references are insufficient, and it is burdensome to perform analysis using these reference genomes. Here, we constructed a set of resources to support NGS analysis using the Japanese reference genome sequence, JG. We created resources for variant calling, gene and repeat element annotations, variant-effect prediction, read mappability, and RNA-seq analysis. We also provide a resource for reference coordinate conversion for further annotation enrichment. We then provide a variant calling protocol using JG-based resources. Our resources provide a guide to prepare sufficient resources for the use of population-specific reference genomes and can facilitate the migration of reference genomes.

## INTRODUCTION

Next-generation sequencing (NGS) studies have identified thousands of genes associated with Mendelian diseases and have supported the diagnosis of many patients worldwide (1, 2). Due to technological advances and significant cost reductions, NGS analyses are now routinely used for basic research in human genetics, cell biological analyses, and as a clinical tool (3, 4).

The reference human genome sequence plays an essential role in NGS analyses (5, 6). In clinical NGS analysis, the raw sequence reads must be analyzed with the appropriate reference genome sequence to prepare for downstream analysis and variant sites can be defined only after comparing the sequence between the reference and the reads from the tested sample. Some studies have shown that different reference sequence yields different results. For example, the difference between using GRCh37 and GRCh38 on germline variant identification amounts to 1.5% and 2.0% in single nucleotide variants (SNVs) and insertions/deletions (indels), respectively (7). In another research, *NOTCH2NLC* repeat expansion associated with neuronal intranuclear inclusion disease was difficult to detect because the region of this gene was assembled erroneously and missed in the human reference genome until the latest version (GRCh38/hg38) (8). Therefore, the selection of the reference sequence requires careful consideration.

The most commonly used reference genome sequences today are GRCh37/hg19 and GRCh38/hg38 (7). They have been maintained and updated by the Genome Reference Consortium (GRC) since their publication in 2009 and 2013, respectively (9, 10). While they are undoubtedly indispensable, some potential problems exist, particularly for some populations. The genetic information used to build the reference genome is biased toward African and European ancestries (11–13). In addition, although multiple donors participated in the project, the reference genome inevitably carried over rare or individual-specific variants, which led to confusion in the interpretations of the variants’ effects (14). Because of these biases, GRC reference genomes may not be ideal for all populations. Accordingly, several population-specific reference genomes have recently been constructed to provide a choice to deal with the vast genetic diversity, such as the Korean reference genome (KOREF) (15), the Chinese reference genome, HX1 (16), another Korean reference genome, AK1 (17), the Swedish reference genomes (18), the Egyptian reference genome (19), the Tibetan reference genome (20), and the Japanese reference genome, JG (21).

Despite the various options for reference genomes, GRC reference genomes are still widely used by most research and clinical laboratories. One of the reasons for the slow migration of references is the lack of supporting resources for analysis with population-specific reference genomes. For GRCh37, much knowledge has been accumulated, and resources are well organized. However, these resources cannot be used for analyses on other reference genomes because the coordinate system or the position of each gene differs. Therefore, it is necessary to transfer information between reference genomes through the so-called “lift-over” process, or the resource should be re-created again, which requires much time and effort.

In this work, we construct a set of resources to support NGS analysis using the Japanese reference genome sequence and provide an example of the resources required for analysis using a population-specific reference genome. In addition, we demonstrated a variant calling procedure with JG using the constructed ecosystem.

## MATERIAL AND METHODS

### Study design

The aim of this study was to improve the environment for NGS data analysis using the Japanese reference genome. To this end, we constructed and provided resources to support NGS studies. These resources are used for variant calling, gene and repeat element annotation, variant-effect prediction, read mappability, and RNA-seq analysis. We also provided information on how to create them so that they can be applied to other references. To validate the use of these resources, we then developed a practical variant calling protocol using JG. The scripts were described in RESULTS for testing and application to other studies. Example data and required resources were described in the pipeline.

### JG2.1.0-based resources for NGS study

We have constructed a set of resources described in Table 1 to support NGS studies with JG2.1.0. First, we created chain files, which are required for lift-over annotations between JG2.1.0 and other genome assemblies. Chain files were created using minimap2 (ver. 2.24) software (22, 23) and transanno (v0.24) software (https://github.com/informationsea/transanno) based on GRCh38 reference sequence. A gene annotation file in GFF3 was lifted over from GRCh38-based GFF3 (https://ftp.ebi.ac.uk/pub/databases/gencode/Gencode_human/release_44/gencode.v44.pri <u>mary_assembly.annotation.gff3.gz</u>) using Liftoff (ver. 1.6) software (24) and was converted to GTF format using GffRead (v0.12.7) software (25). A refFlat format file was created from the GFF3 file using gff3ToGenePred (the UCSC Genome Browser command line tool, https://github.com/ucscGenomeBrowser).

**Table 1.**
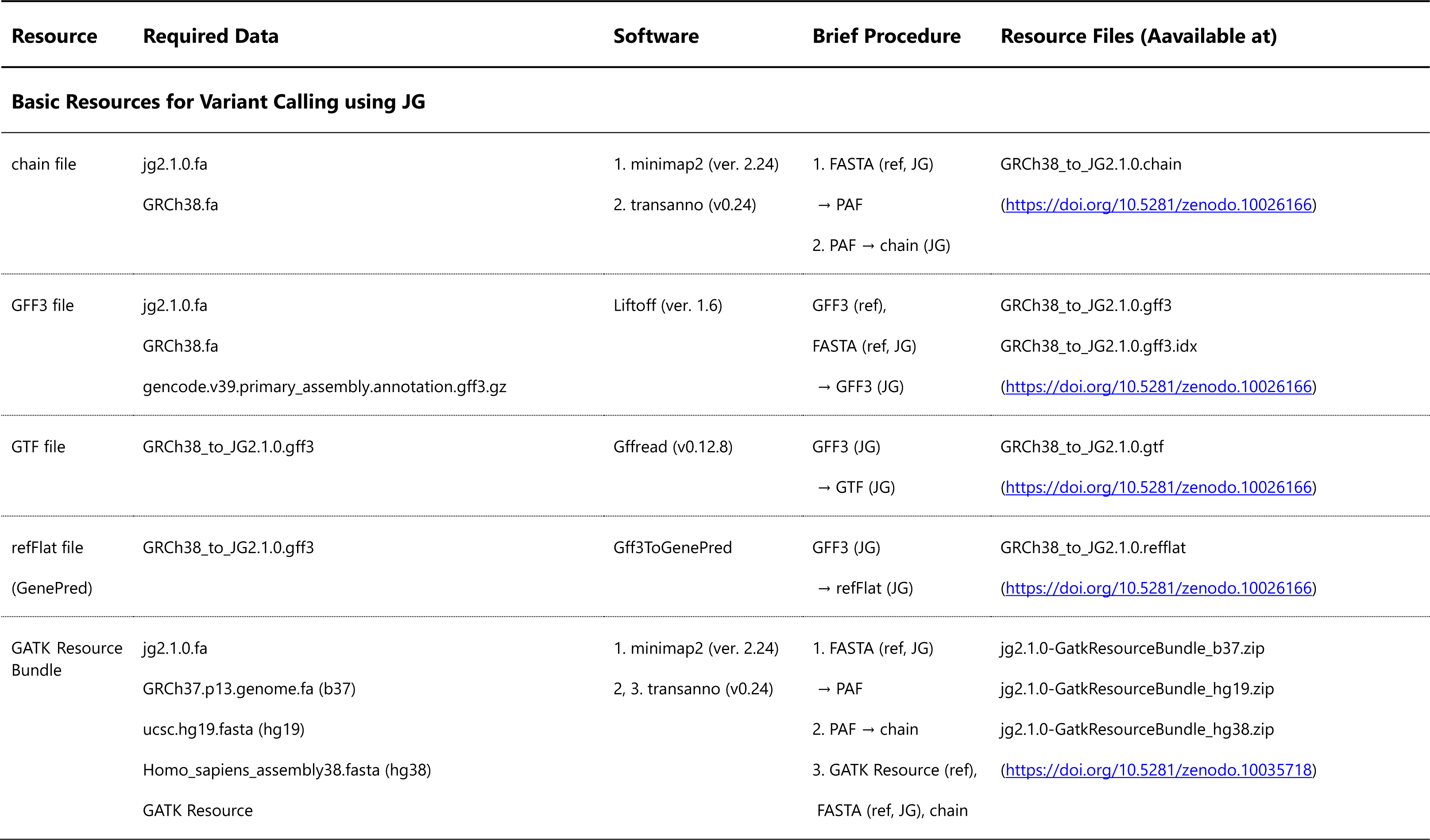
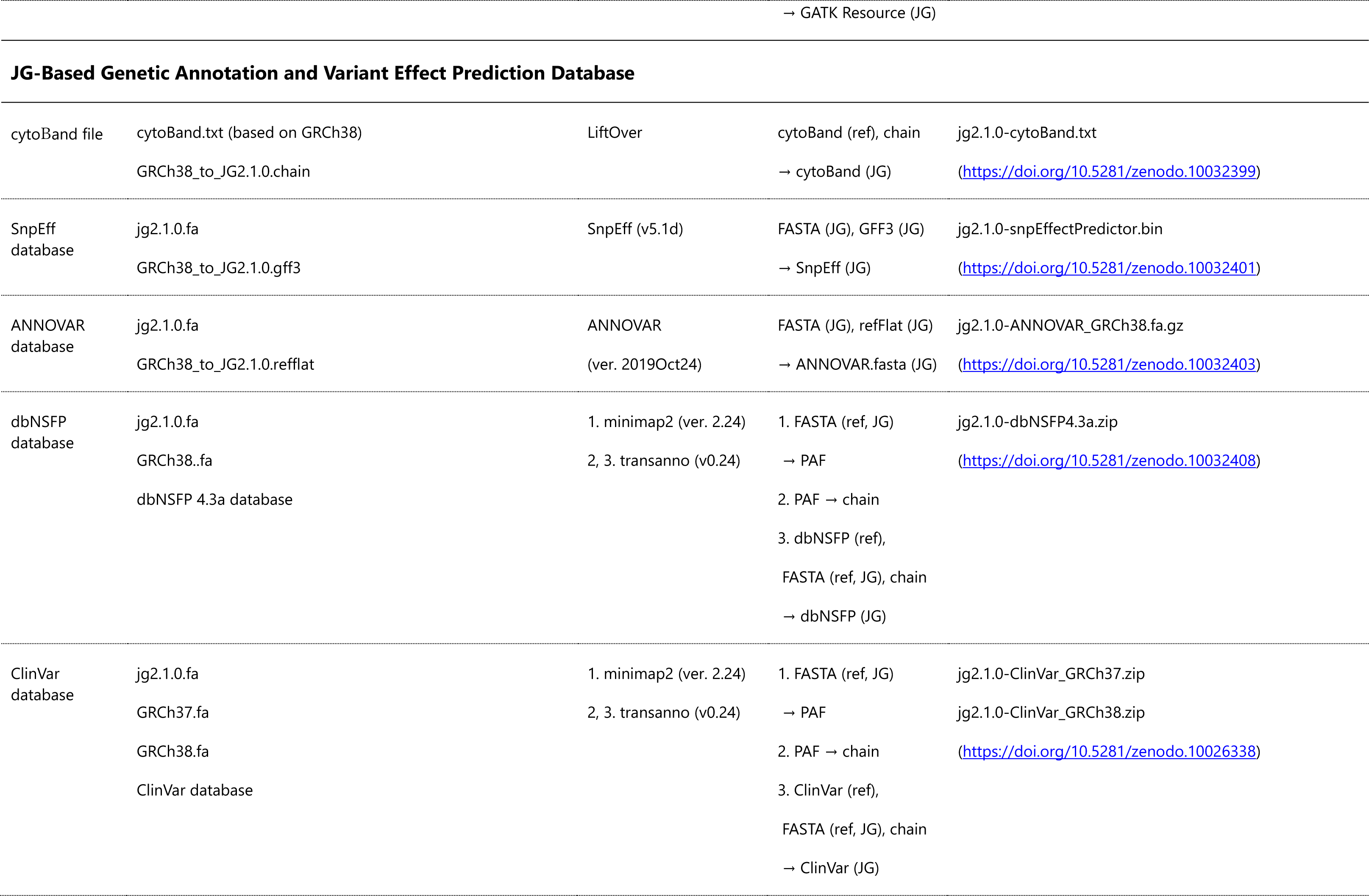
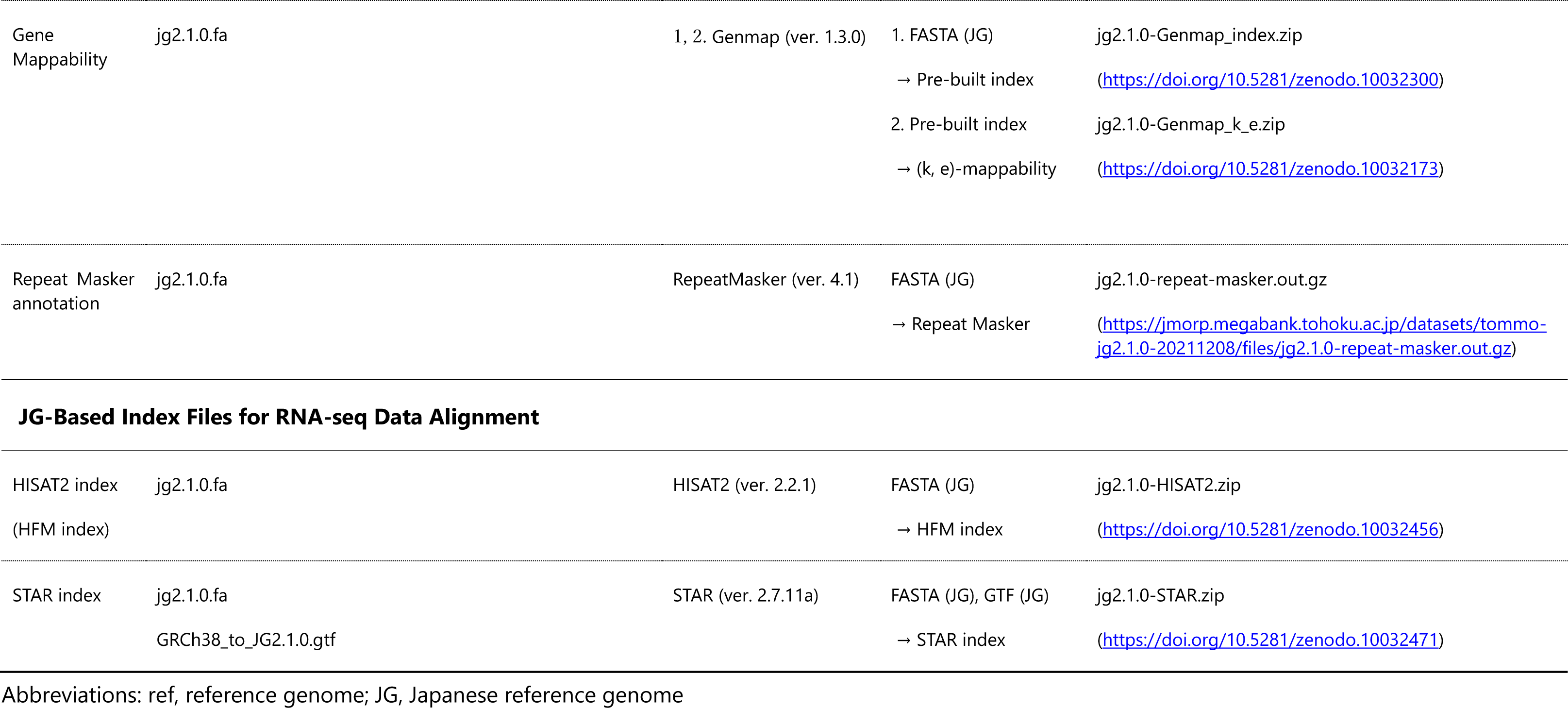
JG2.1.0-Based Resources.

Then, using these files, useful tools and databases for variant analysis were lifted over to JG2.1.0. Genome Analysis ToolKit (GATK) (5) Resource Bundle were created by lifting over using minimap2 and transanno from the corresponding reference genomes. The cytoband file based on GRCh38 (https://hgdownload.soe.ucsc.edu/goldenPath/hg38/database/cytoBand.txt.gz) was lifted over to JG2.1.0 using LiftOver software (26). SnpEff and ANNOVAR databases were created using SnpEff (v5.1d) software (27) and ANNOVAR (ver. 2019Oct24) software (28), respectively. The dbNSFP database (29) was created based on dbNSFP v4.3. The dbNSFP database has two branches: academic (v4.3a) and commercial (v4.3c). While v4.3a includes all the resources, v4.3c does not include several prediction scores (http://database.liulab.science/dbNSFP). The academic version was created here. ClinVar databases were transferred using minimap2 and transanno from GRCh37 and GRCh38, respectively.

In addition, we used RepeatMasker (ver. 4.1) (http://repeatmasker.org) to annotate repeat elements. We created genome mappability resources. Genome mappability, also called (*k*, *e*)-mappability, is the uniqueness of *k*-mers for each position, allowing for up to e mismatches. We calculated the (30,2)-, (36, 0)-, (36, 2)-, (24, 0)-, (24, 2)-, and (50, 2)-mappability of the JG2.1.0 reference sequence using Genmap (ver. 1.3.0) software (30). Besides, we prepared the pre-built index of JG so that one can easily calculate arbitrary (*k*, *e*)-mappability using Genmap. We also built two indices for HISAT2 (ver. 2.2.1) (31) and STAR (ver. 2.7.11a) (32) that are used to align RNA-seq data.

### Variant calling protocol using JG

We developed a variant calling protocol following the GATK Best Practices workflow (6). The test dataset was subsampled from the Han Chinese trio reference samples from the Genome in A Bottle consortium (33) to reduce the computational time and resources. Since paired-end sequencing was used in the analysis, the read data were divided into R1 and R2. We used bwa-mem2 software (34) (ver. 2.2.1) (https://github.com/bwa-mem2/bwa-mem2), SAMtools software (35) (ver. 1.16.1) (https://github.com/samtools/samtools), GATK tools (5) (ver. 4.3.0.0) (https://github.com/broadinstitute/gatk), python 3 software (https://www.python.org/downloads/), SnpEff and SnpShift software (27) (v5.1d) (http://pcingola.github.io/SnpEff/) and Java runtime environment (Java 11 or later) (https://www.oracle.com/java/) in this protocol.

## RESULTS

### provideJG2.1.0-based resources for NGS study

We constructed the resources for short-read sequencing analysis using the Japanese reference genome, JG2.1.0, according to the method described in MATERIALS AND METHODS and Table 1. These resources contain three types of files, depending on their purpose: essential resource files needed for liftover and variant calling, genetic annotation and variant effect prediction database, and index files for RNA-seq data alignment. These resources were provided at the locations listed in Table 1.

### Variant calling process using JG2.1.0-based resources

We validated the variant calling procedure using the JG-based resources we created. In this protocol, we analyzed trio sequencing data following the GATK Best Practices workflow (6). We provided the scripts for this process below.

0. Preparation
  \# Create directories at an arbitrary location $ cd $ mkdir variant_call/ $ mkdir variant_call/materials/ $ mkdir variant_call/intermediate/ $ mkdir variant_call/results/ $ cd variant_call/ # Download the required data and put those files in the appropriate directory The data used in this analysis was obtained from the location described below. We decompressed the downloaded files if necessary and placed them in the appropriate directory (here, variant_call/materials/). # JG2.1.0 FASTA file $ wget -P materials/ https://jmorp.megabank.tohoku.ac.jp/datasets/tommo-jg2.1.0-20211208/files/jg2.1.0.fa.gz $ gzip -cd materials/jg2.1.0.fa.gz > materials/JG.fa && rm materials/jg2.1.0.fa.gz # JG-Based GATK resource bundle $ wget -P materials/ https://jmorp.megabank.tohoku.ac.jp/datasets/tommo-jg2.1.0-20211208/files/JG2.1.0-ResourceBundle-from-b37.zip $ unzip materials/JG2.1.0-ResourceBundle-from-b37.zip -d materials/ && rm materials/JG2.1.0-ResourceBundle-from-b37.zip # Example dataset – subsampled Han Chinese trio 30X reads file (if necessary) $ wget -P materials/ https://zenodo.org/records/7306771/files/HG005_Son.R1.fastq.gz $ wget -P materials/ https://zenodo.org/records/7307245/files/HG005_Son.R2.fastq.gz $ wget -P materials/ https://zenodo.org/records/7307258/files/HG006_Father.R1.fastq.gz $ wget -P materials/ https://zenodo.org/records/7307266/files/HG006_Father.R2.fastq.gz $ wget -P materials/ https://zenodo.org/records/7307277/files/HG007_Mother.R1.fastq.gz $ wget -P materials/ https://zenodo.org/records/7307823/files/HG007_Mother.R2.fastq.gz
1. Alignment The process of creating a bwa-mem2 index used approximately 70 GB of RAM. Output files are available at https://doi.org/10.5281/zenodo.7567194 for the following procedure if it is difficult to perform this process.
  \# create a bwa-mem2-based index of JG for alignment $ cd variant_call/ $ bwa-mem2 index materials/JG.fa # create a fasta.fai file $ samtools faidx materials/JG.fa # create a sequence dictionary of JG $ gatk CreateSequenceDictionary --REFERENCE materials/JG.fa # conduct alignment $ bwa-mem2 mem -R “@RG\tID:son\tSM:son\tPL:Illumina\tLB:son” \
    materials/JG.fa \ materials/HG005_Son.R1.fastq.gz \ materials/HG005_Son.R2.fastq.gz \ > intermediate/son.sam
    $ bwa-mem2 mem -R “@RG\tID:father\tSM:father\tPL:Illumina\tLB:father” \
    materials/JG.fa \ materials/HG006_Father.R1.fastq.gz \ materials/HG006_Father.R2.fastq.gz \ > intermediate/father.sam
    $ bwa-mem2 mem -R “@RG\tID:mother\tSM:mother\tPL:Illumina\tLB:mother” \
    materials/JG.fa \ materials/HG007_Mother.R1.fastq.gz \ materials/HG007_Mother.R2.fastq.gz \ > intermediate/mother.sam
    # convert SAM files to BAM files $ samtools view -bS intermediate/son.sam > intermediate/son.bam $ samtools view -bS intermediate/father.sam > intermediate/father.bam $ samtools view -bS intermediate/mother.sam > intermediate/mother.bam
2. Preprocess for variant calling
  \# sort BAM files by chromosome and coordinate $ samtools sort -o intermediate/son.sort.bam intermediate/son.bam $ samtools sort -o intermediate/father.sort.bam intermediate/father.bam $ samtools sort -o intermediate/mother.sort.bam intermediate/mother.bam # create sorted BAM indexes $ samtools index intermediate/son.sort.bam $ samtools index intermediate/father.sort.bam $ samtools index intermediate/mother.sort.bam # conduct MarkDuplicates process $ gatk MarkDuplicates \
    --INPUT intermediate/son.sort.bam \ --OUTPUT results/son.sort.markdup.bam \ --METRICS_FILE results/son.sort.markdup.metrics.txt \ --CREATE_INDEX TRUE
    $ gatk MarkDuplicates \
    --INPUT intermediate/father.sort.bam \ --OUTPUT results/father.sort.markdup.bam \ --METRICS_FILE results/father.sort.markdup.metrics.txt \ --CREATE_INDEX TRUE
    $ gatk MarkDuplicates \
    --INPUT intermediate/mother.sort.bam \ --OUTPUT results/mother.sort.markdup.bam \ --METRICS_FILE results/mother.sort.markdup.metrics.txt \ --CREATE_INDEX TRUE
3. Base Quality Score Recalibration (BQSR) Although the GATK Best Practices workflow recommends BQSR (6), it is often omitted due to advances in sequencing technologies (36). Because of this, and because we intend to construct original annotations based only on JG, we will skip this process in this protocol and just show the scripts for information.
  \# BQSR $ gatk BaseRecalibrator \
    --reference materials/JG.fa \ --input results/son.sort.markdup.bam \ --known-sites materials/JG2.1.0-ResourceBundle-from- b37/human_g1k_v37_to_JG2.1.0.dbsnp_138.b37.success.sorted.vcf.gz \ --known-sites materials/JG2.1.0-ResourceBundle-from- b37/human_g1k_v37_to_JG2.1.0.Mills_and_1000G_gold_standard.indels.b37.success. sorted.vcf.gz \ --output intermediate/son_recal_data.table
    $ gatk BaseRecalibrator \
    --reference materials/JG.fa \ --input results/father.sort.markdup.bam \ --known-sites materials/JG2.1.0-ResourceBundle-from- b37/human_g1k_v37_to_JG2.1.0.dbsnp_138.b37.success.sorted.vcf.gz \ --known-sites materials/JG2.1.0-ResourceBundle-from- b37/human_g1k_v37_to_JG2.1.0.Mills_and_1000G_gold_standard.indels.b37.success. sorted.vcf.gz \ --output intermediate/father_recal_data.table
    $ gatk BaseRecalibrator \
    --reference materials/JG.fa \ --input results/mother.sort.markdup.bam \ --known-sites materials/JG2.1.0-ResourceBundle-from- b37/human_g1k_v37_to_JG2.1.0.dbsnp_138.b37.success.sorted.vcf.gz \ --known-sites materials/JG2.1.0-ResourceBundle-from- b37/human_g1k_v37_to_JG2.1.0.Mills_and_1000G_gold_standard.indels.b37.success. sorted.vcf.gz \ --output intermediate/mother_recal_data.table
    $ gatk ApplyBQSR \
    --reference materials/JG.fa \ --input results/son.sort.markdup.bam \ --bqsr-recal-file intermediate/son_recal_data.table \ --create-output-bam-index true \ --output results/son.bqsr.bam
    $ gatk ApplyBQSR \
    --reference materials/JG.fa \ --input results/father.sort.markdup.bam \ --bqsr-recal-file intermediate/father_recal_data.table \ --create-output-bam-index true \ --output results/father.bqsr.bam
    $ gatk ApplyBQSR \
    --reference materials/JG.fa \ --input results/mother.sort.markdup.bam \ --bqsr-recal-file intermediate/mother_recal_data.table \ --create-output-bam-index true \ --output results/mother.bqsr.bam
4. Variant calling and Joint calling
  \# conduct variant calling $ gatk --java-options “-Xmx4g” HaplotypeCaller \
    --reference materials/JG.fa \ --emit-ref-confidence GVCF \ --input results/son_sort.markdup.bam \ --output intermediate/son.g.vcf
    $ gatk --java-options “-Xmx4g” HaplotypeCaller \
    --reference materials/JG.fa \ --emit-ref-confidence GVCF \ --input results/father_sort.markdup.bam \ --output intermediate/father.g.vcf
    $ gatk --java-options “-Xmx4g” HaplotypeCaller \
    --reference materials/JG.fa \ --emit-ref-confidence GVCF \ --input results/mother_sort.markdup.bam \ --output intermediate/mother.g.vcf
    # import GVCFs into GenomicsDB by chromosome $ for chr in chr1 chr2 chr3 chr4 chr5 chr6 chr7 chr8 chr9 chr10 chr11 chr12 chr13 chr14 chr15 chr16 chr17 chr18 chr19 chr20 chr21 chr22 chrX chrY chrM do
    gatk --java-options “-Xmx4g” GenomicsDBImport \ --variant intermediate/son.g.vcf \ --variant intermediate/father.g.vcf \ --variant intermediate/mother.g.vcf \ --reference materials/JG.fa \ --genomicsdb-workspace-path intermediate/genomics_database.${chr} \ --intervals ${chr}
    gatk --java-options “-Xmx4g” GenotypeGVCFs \ --reference materials/JG.fa \ --variant gendb://intermediate/genomics_database.${chr} \ --output intermediate/joint_genotyped.${chr}.vcf \ --intervals ${chr}
    done # conduct joint genotyping $ input_files=”” $ for chr in chr1 chr2 chr3 chr4 chr5 chr6 chr7 chr8 chr9 chr10 chr11 chr12 chr13 chr14 chr15 chr16 chr17 chr18 chr19 chr20 chr21 chr22 chrX chrY chrM do
    input_files+=” --INPUT intermediate/joint_genotyped.${chr}.vcf “
    done $ gatk --java-options “-Xmx4G” MergeVcfs \
    --OUTPUT results/merged.vcf.gz \ ${input_files}
5. Filtering - Variant Quality Score Recalibration (VQSR) # create a recalibration model for SNV $ gatk --java-options “-Xmx4g” VariantRecalibrator \
  --reference materials/JG.fa \ --variant results/merged.vcf.gz \ --resource:hapmap,known=false,training=true,truth=true,prior=15.0 materials/JG2.1.0-ResourceBundle-from- b37/human_g1k_v37_to_JG2.1.0.hapmap_3.3.b37.success.modified.vcf \ --resource:omni,known=false,training=true,truth=false,prior=12.0 materials/JG2.1.0-ResourceBundle-from- b37/human_g1k_v37_to_JG2.1.0.1000G_omni2.5.b37.success.sorted.vcf.gz \ --resource:1000G,known=false,training=true,truth=false,prior=10.0 materials/JG2.1.0-ResourceBundle-from- b37/human_g1k_v37_to_JG2.1.0.1000G_phase1.snps.high_confidence.b37.success.sorted.vcf.gz \ --resource:dbsnp,known=true,training=false,truth=false,prior=2.0 materials/JG2.1.0- ResourceBundle-from- b37/human_g1k_v37_to_JG2.1.0.dbsnp_138.b37.success.sorted.vcf.gz \ --use-annotation QD \ --use-annotation MQ \ --use-annotation MQRankSum \ --use-annotation ReadPosRankSum \ --use-annotation FS \ --use-annotation SOR \ --mode SNP \ --max-gaussians 6 \ --output intermediate/snv.recal \ --tranches-file intermediate/snv.tranches
  \# create a recalibration model for INDEL $ gatk --java-options “-Xmx4g” VariantRecalibrator \
  --reference materials/JG.fa \ --variant results/merged.vcf.gz \ --resource:dbsnp,known=true,training=false,truth=false,prior=2.0 materials/JG2.1.0- ResourceBundle-from- b37/human_g1k_v37_to_JG2.1.0.dbsnp_138.b37.success.sorted.vcf.gz \ --resource:mills,known=false,training=true,truth=true,prior=12.0 materials/JG2.1.0- ResourceBundle-from- b37/human_g1k_v37_to_JG2.1.0.Mills_and_1000G_gold_standard.indels.b37.success. sorted.vcf.gz \ --use-annotation QD \ --use-annotation MQ \ --use-annotation MQRankSum \ --use-annotation ReadPosRankSum \ --use-annotation FS \ --use-annotation SOR \ --mode INDEL \ --max-gaussians 4 \ --output intermediate/indel.recal \ --tranches-file intermediate/indel.tranches
  \# conduct VQSR for each of SNV and INDEL recalibration model $ gatk --java-options “-Xmx4g” ApplyVQSR \
  --reference materials/JG.fa \ --variant intermediate/merged.snv.vcf.gz \ --output intermediate/merged.snv.vqsr.vcf.gz \ --tranches-file intermediate/snv.tranches \ --recal-file intermediate/snv.recal \ --create-output-variant-index true \ -mode SNP $ gatk --java-options “-Xmx4g” ApplyVQSR \ --reference materials/JG.fa \ --variant intermediate/merged.indel.vcf.gz \ --output intermediate/merged.indel.vqsr.vcf.gz \ --tranches-file intermediate/indel.tranches \ --recal-file intermediate/indel.recal \ --create-output-variant-index true \ -mode INDEL
  \# merge SNV-filtered VCF and INDEL-filtered VCF $ gatk --java-options “-Xmx4G” MergeVcfs \
  --OUTPUT results/merged.vqsr.vcf.gz \ --INPUT intermediate/merged.snv.vqsr.vcf.gz \ --INPUT intermediate/merged.indel.vqsr.vcf.gz

### VCF Annotation using SnpEff / SnpSift

0. Preparation
  \# download and setup SnpEff $ cd $ mkdir annotation/ $ cd annotation/ $ wget https://snpeff.blob.core.windows.net/versions/snpEff_latest_core.zip $ unzip snpEff_latest_core.zip $ mkdir -p snpEff/data/JG2.1.0/
  \# edit config file We edited the “snpEff.config” file in the SnpEff directory using a text editor. Specifically, we added the following text to the “Non-Standard Databases” section (Fig. 1).

**Figure 1.**
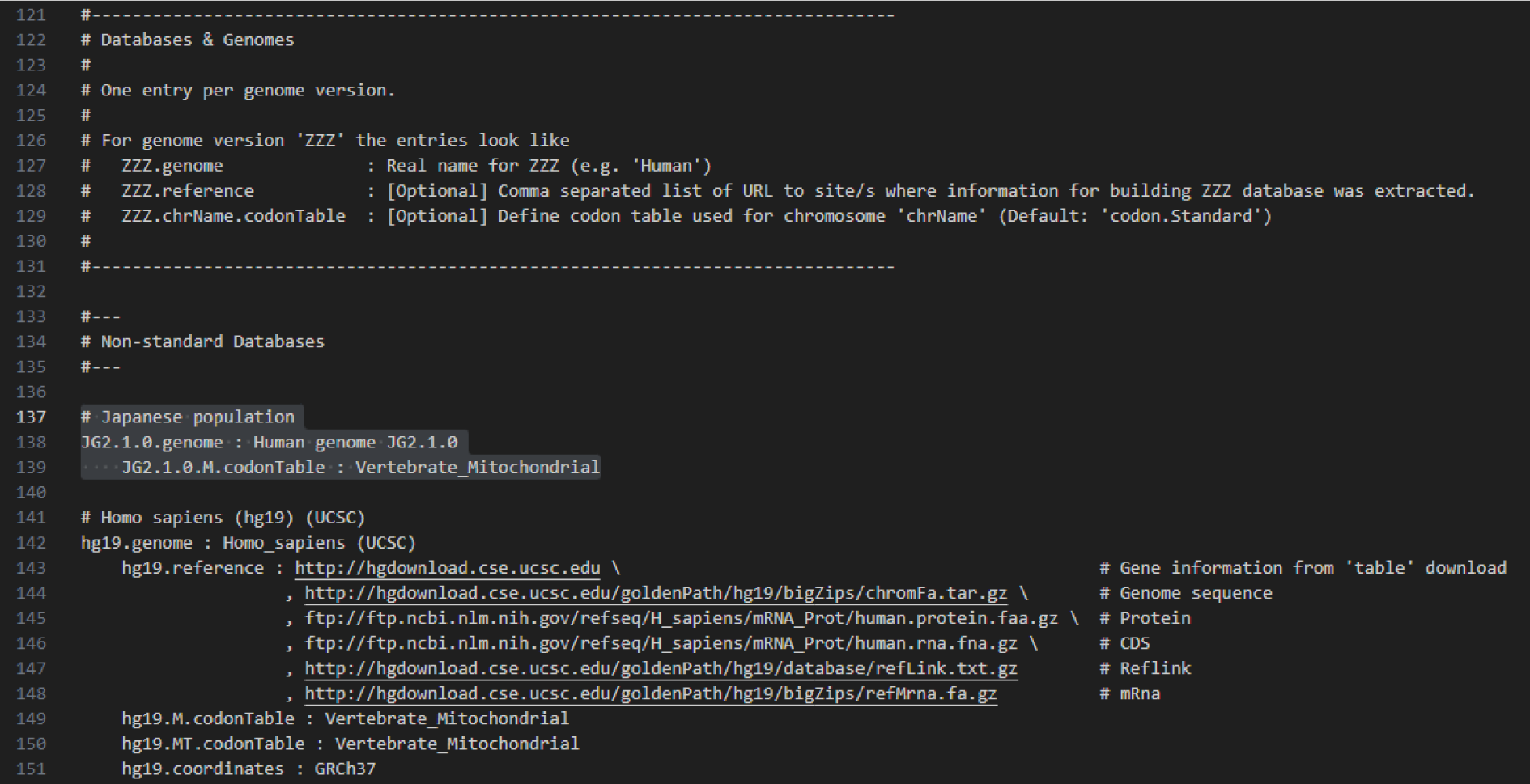
Example of SnpEff.config adjustment. Text for using JG should be added to the “Non-Standard Databases” section. In this example, the text is added to lines 137-139. “# Japanese population JG2.1.0.genome : Human genome JG2.1.0 JG2.1.0.M.codonTable : Vertebrate_Mitochondrial” # data preparation $ cd snpEff/ $ wget -P data/JG2.1.0/ https://zenodo.org/record/7503342/files/snpEffectPredictor.bin $ wget -P data/JG2.1.0/ https://zenodo.org/record/7519222/files/merged.vqsr.vcf.gz $ wget -P data/JG2.1.0/ https://zenodo.org/record/7602190/files/dbNSFP4.3a_variant.chr22.liftover_jg2.1.0.txt.gz $ wget -P data/JG2.1.0/ https://zenodo.org/record/7602190/files/dbNSFP4.3a_variant.chr22.liftover_jg2.1.0.txt.gz.tbi
1. Annotation Using SnpEff
  \# annotate VCF $ java -Xmx8g -jar snpEff.jar \
    JG2.1.0 \ data/JG2.1.0/merged.vqsr.vcf.gz \ > data/JG2.1.0/merged.vqsr.ann.vcf
2. Variant Filtering Using SnpEff # filter out samples with quality less than 35 $ java -jar SnpSift.jar \
  filter “( QUAL >= 35 )” \ data/JG2.1.0/merged.vqsr.ann.vcf \ > data/JG2.1.0/merged.vqsr.ann.filtered.QUAL35.vcf
  \# any variant in chromosome 22 $ java -jar SnpSift.jar \
  filter “( CHROM = ‘chr22’ )” \ data/JG2.1.0/merged.vqsr.ann.vcf \ > data/JG2.1.0/merged.vqsr.ann.filtered.CHROM22.vcf
  \# any transition variant $ java -jar SnpSift.jar \
  filter “(( REF = ‘A’ ) & ( ALT = ‘G’ )) | \
    (( REF = ‘G’ ) & ( ALT = ‘A’ )) | \ (( REF = ‘C’ ) & ( ALT = ‘T’ )) | \ (( REF = ‘T’ ) & ( ALT = ‘C’ ))” \
    data/JG2.1.0/merged.vqsr.ann.vcf \ > data/JG2.1.0/merged.vqsr.ann.filtered.Transition.vcf
  \# any missense variant $ java -jar SnpSift.jar \
  filter “ANN[*].EFFECT has ‘missense variant’” \ data/JG2.1.0/merged.vqsr.ann.vcf \ > data/JG2.1.0/merged.vqsr.ann.filtered.missense.vcf
  \# any variant has high annotation impact $ java -jar SnpSift.jar \
  filter “ANN[*].IMPACT has ‘HIGH’” \ data/JG2.1.0/merged.vqsr.ann.vcf \ > data/JG2.1.0/merged.vqsr.ann.filtered.high_impact.vcf
  \# any variant on TRMT2A gene $ java -Xmx8g -jar SnpSift.jar \
  filter “ANN[*].GENE has ‘TRMT2A’” \ data/JG2.1.0/merged.vqsr.ann.vcf \ > data/JG2.1.0/merged.vqsr.ann.filtered.TRMT2A.vcf
  \# any variant with the gene biotype “protein_coding” $ java -Xmx8g -jar SnpSift.jar \
  filter “ANN[*].BIOTYPE has ‘protein_coding’” \ data/JG2.1.0/merged.vqsr.ann.vcf \ > data/JG2.1.0/merged.vqsr.ann.filtered.protein_coding.vcf
3. SnpSift dbNSFP # annotate using dbNSFP database $ java -jar SnpSift.jar \
  dbnsfp \ -v \ -db data/JG2.1.0/dbNSFP4.3a_variant.chr22.liftover_jg2.1.0.txt.gz \ data/JG2.1.0/merged.vqsr.ann.filtered.CHROM22.vcf \ ­ data/JG2.1.0/merged.vqsr.ann.filtered.CHROM22.dbnsfp.vcf

## DISCUSSION

Here, we constructed the resources for NGS study with the Japanese reference genome sequence, JG2.1.0 (Table 1). And then, we demonstrated the variant calling procedure with trio sequencing data following the GATK Best Practices workflow (6) using these resources. This workflow is expected to yield stable results in NGS studies with the Japanese reference genome. As the technologies have improved and costs have decreased significantly, NGS analysis is increasingly being used for basic research in human genetics and clinical diagnostics (3, 4). Although adequate resources and clear manuals are likely to be required, in practice, except for JG, such an ecosystem is not yet sufficient.

Recently, several population-specific genome assemblies have been constructed to deal with the vast genetic diversity (15–21). The use of these references, adjusted for population specificity, is expected to provide an advantage in NGS analysis, e.g., efficient variant identification in clinical practice. Notably, a case of a Japanese female in whom a breakpoint of reciprocal translocations was detected only in JG was reported (37). On the other hand, the existence of multiple references has been criticized as complicating the evaluation of the analysis (38). In addition, many attempts to construct population-specific reference only provide the assembled DNA sequences in FASTA format, so that we must assign contigs to chromosomes. Furthermore, we need annotations and various analytical resources.

With the recent advent of long-read technologies, the Telomere-to-Telomere (T2T) sequence has been constructed (39). The T2T-CHM13 achieved a near gapless telomere-to-telomere assemblies. Although the T2T-CHM13 reference will be the next prevailing reference for human genetics (40), the T2T-CHM13 genome was derived from only one European cell line and hence has a population bias (39). Another new attempt to address the limitations of current references is a genome graph (41, 42). The genome graph aims to comprehensively assess human genetic variation by including the vast genetic diversity using graph representation (41). The graph representation may replace the conventional references in the near future (41–43), but conventional linear reference genomes will continue to be used due to their high affinity with previous analyses and ease of interpretation at least in some field. However, when using a linear population-specific genome reference, a significant number of accompanying resources must be prepared for analysis, which requires a certain amount of knowledge and effort. Therefore, we constructed the resources necessary to analyze short-read sequencing using JG2.1.0.

There are several limitations in this work. First, the variant calling protocol we described in this paper is for JG only and it assumes that supporting resources are available. And second, the resources we constructed are JG-specific. Therefore, similar resources are required for migration to another reference of interest. However, we also provided how to create these resources. Our protocol will allow the development of resources to support NGS analysis and enable its application to other references.

Today, with the widespread use of NGS analysis, population-specific reference is one of the choices to deal with the vast genetic diversity. However, references alone are not sufficient to handle the burden of migrating references and running analyses, so we have created a set of resources based on JG and provided them here. The development of an ecosystem for each reference is an important foundation giving users “analysis ready”.

## DATA AVAILABILITY

All the resources we have constructed are located at the repository sites listed in Table 1.

## AUTHOR CONTRIBUTIONS

Tomohisa Suzuki: Methodology, Resources, Writing—original draft. Kota Ninomiya: Methodology, Resources, Writing— review & editing. Takamitsu Funayama: Methodology, Resources, Writing— review & editing. Yasunobu Okamura: Resources, Writing— review & editing. Shu Tadaka: Resources, Writing— review & editing. Kengo Kinoshita: Resources, Writing— review & editing. Masayuki Yamamoto: Conceptualization, Writing— review & editing. Shigeo Kure: Supervision, Writing— review & editing. Atsuo Kikuchi: Supervision, Writing— review & editing. Gen Tamiya: Conceptualization, Supervision, Writing— review & editing. Jun Takayama Conceptualization, Methodology, Project administration, Writing— review & editing.

## ACKNOWLEDGEMENTS

We express our gratitude to Dr. Justin Matthew Zook and the Genome in a Bottle Consortium for granting us permission to use and distribute subsampled Han Chinese trio genomic data. We also thank Dr. Xiaoming Liu for allowing us to publish the JG-based dbNSFP database. Finally, we appreciate all the volunteers who participated in the Tohoku Medical Megabank project.

